# Genomic insights into the virulence repertoire and hemibiotrophic lifestyle of the grapevine black rot pathogen *Phyllosticta ampelicida*

**DOI:** 10.1101/2025.05.08.652932

**Authors:** Monica Colombo, Paola Bettinelli, Jadran Garcia, Giuliana Maddalena, Silvia Laura Toffolatti, Ludger Hausmann, Silvia Vezzulli, Simona Masiero, Dario Cantù

**Affiliations:** Council for Agricultural Research and Economics - Research Centre for Genomics and Bioinformatics, Fiorenzuola d’Arda, PC, Italy; Grapevine Physiology and Breeding Unit, Research and Innovation Centre, Fondazione Edmund Mach, 38098, San Michele all’Adige, TN, Italy; Department of Viticulture and Enology, University of California, Davis, CA, 95616, United States; Dipartimento di Scienze Agrarie e Ambientali, Università degli Studi di Milano, 20133, Milano, MI, Italy; Julius Kuhn Institute - Institute for Grapevine Breeding Geilweilerhof, Siebeldingen, Germany; Dipartimento di Bioscienze, Università degli Studi di Milano, Milano, MI, Italy; Genome Center, University of California, Davis, Davis, CA, 95616, United States

**Keywords:** fungal pathogen virulence factors, pathogenomics, cell wall degrading enzymes, secondary metabolism, effectors, *Guignardia bidwellii*, *Vitis vinifera*

## Abstract

*Phyllosticta ampelicida*, the causal agent of grapevine black rot, is a globally emerging pathogen that infects all grapevine green tissues, with young shoots and berries being particularly susceptible. Severe infections can result in total crop loss. To investigate its virulence repertoire, we generated a high-quality genome assembly of strain GW18.1 using long-read sequencing, resulting in 22 scaffolds, including four complete chromosomes and 12 chromosome arms, with a total genome size of 35.6 Mb and 10,289 predicted protein-coding genes. Two additional strains (TN2 and LB22.1) were sequenced with short reads to assess intraspecies diversity. Comparative genomics revealed a conserved virulence factor repertoire, including 314 carbohydrate-active enzymes (CAZymes), 17 cytochrome P450s, 35 peroxidases, and 20 secondary metabolite biosynthetic gene clusters (BGCs). Trophic lifestyle prediction based on gene content supports a biotrophic-like lifestyle consistent with hemibiotrophic pathogens. Broader comparisons with other *Phyllosticta* species and ten plant-pathogenic fungi pointed to species-specific features, while analysis of gene family evolution identified expansions and contractions in transporters and CAZymes. These genomic resources will support efforts to better understand and manage grapevine black rot.

## Introduction

Plant diseases cause 10–15% of global crop losses annually, amounting to hundreds of billions of dollars (Chatterjee et al. 2016). Fungi, responsible for 70–80% of these diseases, are a major threat to sustainable agriculture (Marín-Menguiano et al. 2019; Stukenbrock and Gurr 2023). Grapevines depend heavily on fungicides, accounting for over 65% of all agricultural fungicide use, a figure expected to rise (Nagesh et al. 2023). Black rot, a key disease in temperate-humid regions along with downy and powdery mildew, originated in North America and was introduced to Europe in 1885 (Pirrello et al. 2019). It spread rapidly due to the susceptibility of European grapevines and is now found worldwide, likely due to the movement of infected plant material (Pirrello et al. 2019).

Grapevine black rot is caused by the fungus *Phyllosticta ampelicida* [Engleman] Van der Aa (syn. *Guignardia bidwellii* (Ellis) Viala and Ravaz), a member of the class dothideomycetes, order Botryosphaeriales, family Botryosphaeriaceae, and genus *Guignardia* (**Figure 1A**). The pathogen infects all green grapevine organs and causes significant damage. *Phyllosticta ampelicida* is classified as a hemibiotrophic ascomycete, characterized by an initial biotrophic, asymptomatic phase followed by a necrotrophic phase associated with visible symptoms and tissue damage. On leaves, infection presents as circular lesions that become brown with dark-reddish borders; pycnidia (asexual fruiting bodies) appear within the central necrotic area (**Figure 1B**). On young shoots, the pathogen produces dark, elongated spots on the first internode, which may expand, girdle the shoot, and penetrate the tissue, leading to cracking or canker formation. Black pycnidia are also commonly observed at the center of these lesions. The most characteristic symptoms occur on berries, from fruit set to late veraison, beginning as small, light-brown spots that expand across the fruit surface (**Figure 1C**). Infected berries soften, become spongy, then dry out into blackish-blue mummies, often bearing visible pycnidia (Kuo and Hoch 1996b). Under favorable environmental conditions, *Phy. ampelicida* can cause severe yield losses. Even low disease severity can significantly impact production due to the non-linear relationship between disease intensity and yield loss (Molitor and Beyer 2014).

**Figure 1.**
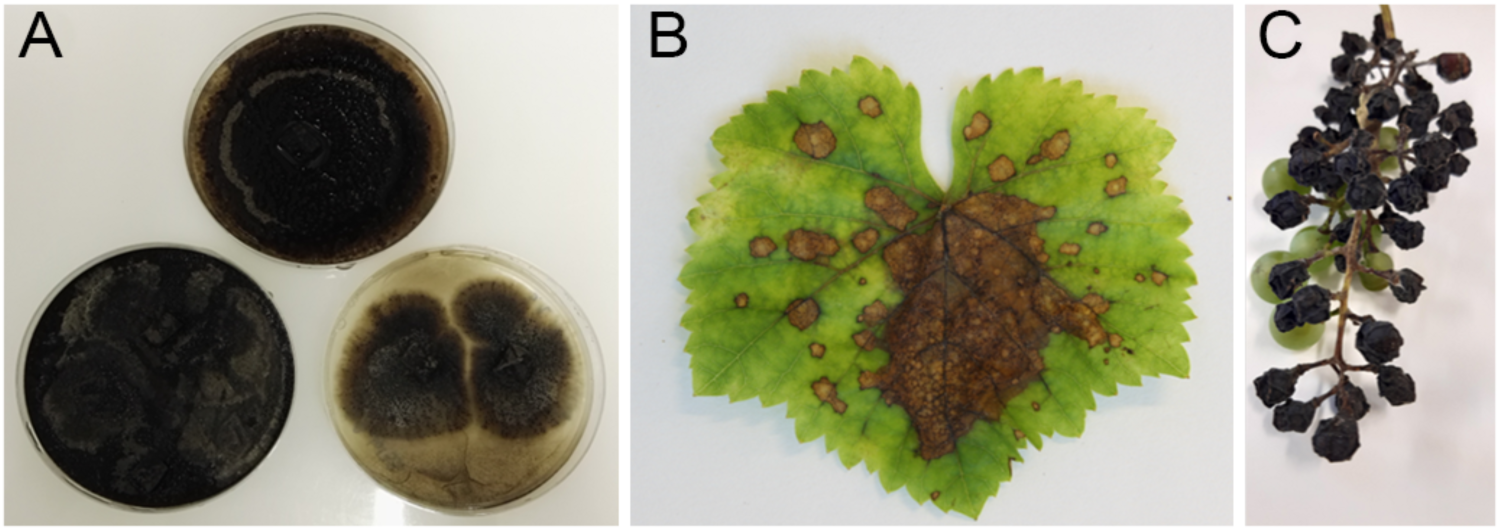
Grapevine black rot. **A**) Axenic culture of the causal agent *Phy. ampelicida* (top TN2, bottom left GW18.1, bottom right LB22.1). Disease symptoms on grapevine **B**) leaves and **C**) fruits.

The rising incidence of black rot is linked to reduced fungicide use due to the adoption of disease-resistant grapevine varieties and increased mechanical harvesting and pruning, which fail to remove infected plant material (Ullrich et al. 2009; Molitor and Beyer 2014; Hausmann et al. 2017). Mummified grape bunches often remain on vines over winter, releasing ascospores in spring to initiate primary infections, while pycnidiospores spread the disease during the growing season, with both spore types germinating similarly (Ullrich et al. 2009). In Europe, the black rot surge also coincides with the ban of sterol biosynthesis inhibitors. Additionally, abandoned and uncultivated vineyards in Central Europe exacerbate the problem by acting as reservoirs, with infectious material easily spread by wind to nearby cultivated areas.

To address the rise of plant pathogen resistance and reduce reliance on pesticides, alternative disease management strategies are needed. The identification of genetic resistance loci and associated molecular markers has enabled marker-assisted breeding and gene stacking in grapevine improvement programs (Vezzulli et al., 2019). More recently, genome-guided studies have begun to identify the causal genes underlying resistance QTLs, opening new opportunities for targeted genome editing (Cantu et al. 2024). In parallel, the analysis of plant pathogen genomes is critical for understanding disease emergence and spread, supporting the development of advanced diagnostics and effective curative or preventive measures (Paineau et al. 2024). New plant protection products that target essential pathogen proteins and the molecular networks activated during early infection stages are also promising (Colombo et al. 2020; Rosa et al. 2023).

A deeper understanding of pathogen virulence mechanisms is essential to guide these efforts (Paineau et al. 2024). Fungal virulence factors, such as carbohydrate-active enzymes, cytochrome P450s, peroxidases, transporters, secondary metabolites, and effectors play central roles in host colonization and symptom development. These factors support key processes such as cell wall penetration, nutrient acquisition, suppression of host defenses, and induction of tissue necrosis (Horbach et al. 2011; Li et al. 2025). For example, pathogens breach plant cell walls through mechanical pressure and cell wall– degrading enzymes (Pryce-Jones et al. 1999), and counteract oxidative stress via reactive oxygen species (ROS)–scavenging mechanisms such as peroxidases (Mir et al. 2015). Fungal secondary metabolites released during early infection may weaken plant defenses without triggering cell death, promoting pathogen success during the biotrophic phase (van Doorn et al. 2011; Reveglia et al. 2022).

Building on the recent release of a draft genome for *Phy. ampelicida* based on short-read sequencing (Eichmeier et al. 2022), we conducted further genome sequencing using long-read sequencing of a strain isolated in Germany. Additionally, to begin exploring the genetic variation within the species we sequenced two strains isolated in northern Italy using short-read sequencing. To provide a broader perspective on the *Phy. ampelicida* virulence repertoire, comparative genomic analyses were performed within the *Phyllosticta* genus and across grapevine-pathogenic ascomycetes.

## Material and methods

### Isolation of *Phyllosticta ampelicida* gDNA for SMRT and short-read sequencing

A strain of *Phy. ampelicida* isolated by Fondazione Edmund Mach (FEM, San Michele all’Adige, Italy) in Trentino (Italy) (TN2), and one isolated by the Julius Kuhn Institute (JKI)–Institute for Grapevine Breeding Geilweilerhof (Siebeldingen, Germany) (GW18.1), were previously genetically characterized (Bettinelli et al. 2023b) (**Figure 1**). These fungal cultures were maintained at 24° on organic oatmeal agar medium (0.5% w/v) (Bettinelli et al. 2023b). To confirm the purity of the fungal isolates before sequencing, the DNA extracted was subjected to specific diagnostic PCRs followed by sequencing. The samples were confirmed as *Phy. ampelicida* by ITS analysis using primers ITS4 (TCCTCCGCTTATTGATATGC) and ITS5 (GGAAGTAAAAGTCGTAACAAGG) (Rinaldi et al. 2017).

The *Phy. ampelicida* strain LB22.1 was isolated from infected leaves during the 2022 growing season in Traona, Sondrio province, Lombardy (Italy) and maintained on potato dextrose agar (PDA) plates at 22°C. Identification as *Phy. ampelicida* was confirmed through ITS analysis using primers ITS1F (TCCGTAGGTGAACCTGCGG) and ITS4R (TCCTCCGCTTATTGATATGC) (Toffolatti et al. 2007; Horváth et al. 2024). High molecular weight genomic DNA (gDNA) of *Phy. ampelicida* isolates was extracted following the protocol of Morales-Cruz et al. (2020) with minor modifications. The quality of the extracted gDNA was assessed using a NanoDrop UV/Vis spectrophotometer and 0.8% (w/v) agarose gel. The gDNA was quantified with a Qubit 3.0 fluorometer using a Qubit dsDNA BR Assay Kit (Thermo Fisher Scientific, USA).

### Library preparation and genome sequencing

The genome of *Phy. ampelicida* GW18.1 strain was subjected to PacBio CLR Single Molecule Real-time sequencing Technology (SMRT) in order to obtain high-quality long reads genomic data. To ensure optimal sequencing results, gDNA was first cleaned using 0.45x AMPure PB beads (Pacific Biosciences, Menlo Park, CA, USA) prior to library preparation. The SMRTbell template was then prepared using the SMRTbell Express Template Preparation kit 2.0 (Pacific Biosciences, Menlo Park, CA, USA), following the manufacturer’s instructions, with 12 µg of sheared DNA. The SMRTbell template was size-selected with a cutoff size of 17-80 kbp using the Blue Pippin instrument (Sage Science, Beverly, MA, USA), and the resulting size-selected library was cleaned using 1x AMPure PB beads. Finally, the library was sequenced on a SMRT cell using the PacBio Sequel II platform at the DNA Technology Core Facility of the University of California, Davis (CA-USA).

### Genome assembly and scaffolding

PacBio sequencing reads for *Phy. ampelicida* were assembled using Falcon assembler (v.2017.06.28-18.01; Chin et al. 2016). Multiple parameter combinations were tested to achieve the lowest fragmentation. Repetitive content identification in both raw and error-corrected reads was employed, as described in Minio et al. (2019). The resulting contigs were then polished using Arrow from v.GCpp 1.0.0-cd36561 (https://github.com/PacificBiosciences/gcpp). Scaffolding was performed using SSPACE-longreads v.1.0 (Boetzer and Pirovano 2014). To assess the assembly completeness, we performed Benchmarking Universal Single-Copy Orthologs (BUSCO v5.4.2; Manni et al. 2021) analysis with the fungi_odb10 lineage dataset. Illumina sequencing reads from TN2 and LB22.1 were quality-filtered and adapter clipped using Trimmomatic v.0.36 (Bolger et al. 2014), with the following settings: LEADING:7, TRAILING:7, SLIDINGWINDOW:10:20, and MINLEN:140. SPAdes v.3.13.0 (Bankevich et al. 2012) was used to assemble the quality-filtered reads with the careful option and automatic read coverage cutoff after optimizing the multiple Kmer combination. To assess the assembly completeness of the genomes, we performed Benchmarking Universal Single-Copy Orthologs (BUSCO v.5.4.2; Manni et al. 2021) analysis with the fungi_odb10 lineage dataset. Additionally, the package tidk v.0.2.63 (Brown et al. 2025) was used with the search mode to identify the telomeric repeat “CCCTAA” in 1kb windows.

### Variant analysis

Quality-filtered reads from the isolates TN2 and LB22.1 were mapped in a paired-end mode to the genome of the isolate GW18.1 using default parameters in BWA v.0.7.17 (Li and Durbin 2009). PCR and optical duplicates were removed with Picard tools v.2.0.1 (http://broadinstitute.github.io/picard/). HaplotypeCaller (GATK v.4.0.12.0; McKenna et al. 2010) was used to call sequence variants between the isolates using the parameters --ploidy 1 --min_base_quality_score 20. The variants were normalized using BCFtools norm v.1.9-94-g9589876 (Danecek et al. 2021) using the option -m-any. The SNP and INDEL variants up to 50bp were extracted using BCFtools view. For larger variants (larger than 50bp) the genomes of the two isolates were aligned to the reference GW18.1 using Minimap2 v.2.12 (https://academic.oup.com/bioinformatics/article/34/18/3094/4994778?login=true) using the parameters -a -x asm5 --cs. The alignments were converted to bam format, and the tool SVIM-asm v.1.0.3 (https://academic.oup.com/bioinformatics/article/36/22-23/5519/6042701) was used in the haploid mode to annotate INDELS larger than 50bp. The functional impact of SNPs and INDELS was predicted with SnpEff v.5.1 (Cingolani et al. 2012) using default parameters.

### Functional annotation and trophic lifestyle inference

Repeat model libraries were predicted for *Phy. ampelicida* using RepeatModeler v.open-1.0.11 (Smit et al. 2015b) with default parameters. To mask the identified repeats RepeatMasker v.open-4.0.6. (Smit et al. 2015a) was employed along with the known fungal model libraries in the RepBase (v.20160829). For gene model prediction BRAKER1 was used with the option --fungus (Hoff et al. 2016). The Augustus v.3.2.1 (Stanke et al. 2006) version was used to train the models based on the alignment of RNAseq data on the assembled genome. The HISAT2 v2.1.0 (Kim et al. 2019) tool with the very-sensitive option was employed to align RNA-seq data on the assembled genome. The alignment was sorted and prepared for BRAKER1 using samtools-1.3.1 (Li et al. 2009). The predicted proteins were annotated based on the similarity to conserved domains in the Pfam database (Finn et al. 2016). For functional annotation, various databases and parameters (**Supplementary Table 1**) were used. CAZymes were annotated using dbCAN3 (Zheng et al. 2023), keeping only genes annotated with at least two of the three algorithms. Signal peptides were identified using SignalP 5.0 (Almagro Armenteros et al. 2019), and proteins with annotation in both SignalP5 and dbCAN3 databases were annotated as secreted CAZymes. Secondary metabolite clusters were annotated using antiSMASH 6.0 (Blin et al. 2021), while peroxidases were annotated using a specialized database for fungi, fPoxDB (Choi et al. 2014). Cytochrome P450 proteins were annotated using CYPED 6.0 (Sirim et al. 2009). Next, proteins involved in transportation functions were annotated using TCDB (Saier et al. 2006, 2016). Finally, proteins with signal peptides were analyzed for transmembrane domains with TMHMM 2.0 (Krogh et al. 2001). Proteins with no predicted transmembrane domain (TM) in the first 60 amino acids or no more than two TMs in total were used as input for effectorP3 (Han et al. 2022). These were used to annotate effector proteins as apoplastic, cytoplasmic or with dual localization based on the probability values obtained with the software. If the difference between the probabilities was lower than 0.1, the effector protein was annotated as dual localization. The trophic lifestyle prediction of the organisms in this analysis was performed using CATASThrophy (Hane et al. 2020) with default parameters using the predicted protein of the species.

### RNA-seq analysis

Total RNA extraction from *Phy. ampelicida* GW18.1 *in vitro* culture, along with inoculated and mock-inoculated leaves (Bettinelli et al. 2023a) of *V. vinifera* cultivar ‘Pinot noir’, was performed according to the CTAB protocol of Amrine et al. (2015). Stranded mRNA sequencing libraries (KAPA Stranded mRNA-Seq Kit. Roche, Switzerland), quality control and quantification were performed at the Sequencing and Genotyping Platform of Fondazione Edmund Mach (San Michele all’Adige, Italy). The sequencing was carried out on an Illumina Novaseq 6000 platform (Illumina, CA-USA) with paired-end runs of 2 × 150 bps at CIBIO Sequencing platform (Trento, Italy). RNA-seq reads were quality-filtered and adapter-clipped using Trimmomatic v.0.36 (Bolger et al. 2014), with the following settings: LEADING:7. TRAILING:7. SLIDINGWINDOW:10:20. and MINLEN:36. Quality-trimmed reads from the pure fungi were then mapped to the *Phy. ampelicida* genome assembly. For the grapevine inoculated and mock-inoculated samples, reads were mapped to a combined reference of the *V. vinifera* (VITVvi_vPinNoir123_v1.0) and *Phy. ampelicida* genomes using HISAT2 v2.1.0 (Kim et al. 2019) with the very-sensitive option. Alignment files were quantified using Salmon v1.5.1 (Patro et al. 2017) with 100 bootstraps and seqBias and posBias options.

### Comparative genomics

Genomes and gene annotations from the four *Phy. ampelicida* isolates (GW18.1, TN2, LB22.1, PA1) were compared to those of ascomycete species listed in **Supplementary Table 2**. Several *Phyllosticta* species with different levels of pathogenicity were included in the analysis. Additionally, eleven other species responsible for different plant diseases were used for functional comparison. Finally, the basidiomycetes *Fomitiporia mediterranea* and *Stereum hirsutum* along with the non-pathogenic species *Saccharomyces cerevisiae* were used as outgroups. The predicted proteins of all the genomes were used as input for OrthoFinder v.2.5.4 using default parameters. The Single Copy Orthologs obtained were aligned using MUSCLE v.5.1 (Edgar 2004) with the option “-maxiters 16”. The concatenated alignments were parsed with Gblocks v.0.91b (Castresana 2000). The parsed alignments were used to optimize the evolutionary model using ModelTest-NG v.0.1.7 (Darriba et al. 2020). The maximum likelihood tree of the species was created with RAxML-NG v.0.9.0 (Kozlov et al. 2019), using the parsed alignment, and the optimized evolutionary model “LG+I+G4+F” with the options “--tree pars{10} --bs-trees 100”. The clock-calibrated tree was constructed using BEAST v.2.7.6 (Bouckaert et al. 2019). The parsed alignment was prepared with BEAUti v2.7.6 (Bouckaert et al. 2019). Calibration points were set for the ascomycetes crown to 588 million years ago (Mya) (Beimforde et al. 2014) with a normal distribution, and the dothideomycetes group set to 350 Mya (Beimforde et al. 2014) with a normal distribution. Six independent Markov chain Monte Carlo runs of 1,000,000 generations were applied. The LG substitution model with four gamma categories, and the Birth-Death model was used. Tree sampling was done every 1,000 generations. LogCombiner v.2.7.6 (Bouckaert et al. 2019) was used to combine the resulting log and tree files. TreeAnnotator v2.7.6 (Bouckaert et al. 2019) was used to generate the maximum clade credibility tree using a burn-in of 10,000 generations. Figtree (Rambaut 2018) was used to plot and annotate the phylogenetic trees.

### Gene family expansion and contraction analysis

The predicted proteins in all the genomes were clustered following the methods described in Garcia et al. (2024a). The resulting file was prepared for CAFE using the script cafetutorial_mcl2rawcafe.py at https://github.com/hahnlab/cafe_tutorial/tree/main/python_scripts. The resulting file was used to run CAFE with the option “-P 0.0100”, an estimated lambda value of 0.00063887950005574, and the clock-calibrated tree. Families with significant rates of gain or loss of genes (P value < 0.01) were extracted. CafePlotter (https://github.com/moshi4/CafePlotter) was used to annotate the clock-calibrated tree with the number of families expanding and contracting. A Fisher’s exact test was used to obtain the functions that were significantly enriched in the expanded and contracted families of the species of interest.

## Results and discussion

### A highly contiguous *Phyllosticta ampelicida* genome assembly

PacBio CLR sequences of *Phy. ampelicida* GW18.1 were *de novo* assembled into 22 scaffolds with an N50 of 1.9 Mb and a BUSCO completeness score of 99.3%, confirming the completeness of the assembly. Four of the scaffolds were complete chromosomes and 12 were chromosome arms based on the presence of telomeric repeats (**Supplementary Table 3**). The genome spans a total of 35.6 Mb (**Table 1**) that is about 16.7% larger than the draft genome reported by Eichmeier et al. (30.5 Mb, Eichmeier et al. 2022) and the largest *Phyllosticta* genome sequenced to date (Guarnaccia et al. 2019; Rodrigues et al. 2019; van Ingen-Buijs et al. 2024). The increase in size is likely due to the use of long-read sequencing, which improves contiguity and captures repetitive regions more effectively than short-read approaches (Chin et al. 2016). Supporting this, repetitive sequences accounted for 23% of the genome (**Supplementary Table 4**), much higher than in other grapevine pathogens (e.g., 5% for *Neofusicoccum parvum*; Garcia et al. 2024b). Of the 8 Mb of repeats, 92% were interspersed, with 87% unclassified and 13% identified as transposable elements (TEs), dominated by long terminal repeats (LTRs, 74%). Short tandem repeats (STRs) comprised the remaining 8%. Gene prediction identified 10,289 coding DNA sequences (CDSs), closely matching the 10,691 previously reported for this species (Eichmeier et al. 2022). Of these, nearly 80% (8,115) matched annotated genes, with 75% (7,704) carrying complete Pfam domains (**Table 2**). Additionally, signal peptide predictions indicate that about 7% of the proteins may be secreted.

**Table 1.** Summary of genome sequencing statistics of *Phy. ampelicida* isolates.

|  | GW18.1 | LB22.1 | TN2 | PA1* |
| --- | --- | --- | --- | --- |
| Sequencing technology | PacBio CLR | Illumina NovaSeq | Illumina NovaSeq | Illumina MiniSeq |
| Genome size (Mb) | 35.6 | 33.1 | 33.1 | 30.55 |
| Scaffolds | 22 | 11,063 | 10,944 | 6,675 |
| N50 Length | 1.9 Mbp | 90.2 kbp | 85.3 kbp | 20.6 kbp |
| N90 Index (or L90) | 15 | 549 | 580 | 1.768 |
| BUSCO completeness (**) | 99.3% | 99.2% | 99.3% | 98.8% |
| Repeat content (Mbp) | 8.0 (22.6%) | 5.4 (16.4%) | 5.4 (16.4%) | 3.1 (10.2%) |
| N. CDSs | 10,289 | 8,804 | 8,802 | 7,983 (***) |
| Mean protein size (AA) | 543 | 505 | 505 | 509 (***) |

(\*) Published in Eichmeier et al. 2022. (\*\*) Percentage of complete BUSCO peptides found in the assembled genomes. (\*\*\*) Numbers calculated based on a new gene annotation made in this work for the previously published genome.

**Table 2.** Summary of predicted protein-coding genes in *Phy. ampelicida* GW18.1, categorized by annotated functions and number of genes affected by high-impact variants.

| Category | Total |  | Affected by high impact variants |  |
| --- | --- | --- | --- | --- |
|  | Count | % | Count | % |
| Total genes | 10,289 |  | 354 | 3.40% |
| Genes without functional annotation | 2,196 | 21.34 | 125 | 35.31% |
| Genes with functional annotation | 8,093 | 78.66 | 229 | 64.69% |
| PFAM | 7,704 | 74.88 | 212 | 2.80% |
| Transporters | 2,190 | 21.28 | 25 | 1.10% |
| Cytochrome P450 proteins | 54 | 0.52 | 2 | 3.70% |
| Signal peptides | 718 | 6.98 | 18 | 2.50% |
| CAZymes | 314 | 3.05 | 5 | 1.60% |
| Secreted CAZymes | 169 | 1.64 | 2 | 1.20% |
| Secondary metabolites involved genes | 277 | 2.69 | 7 | 2.50% |
| Peroxidases | 35 | 0.34 | 1 | 2.90% |

### Sequence and structural variation across *Phyllosticta ampelicida* genomes

Differences in *Phy. ampelicida* strain virulence may contribute to varying susceptibility levels of the same grapevine varieties to black rot (Bettinelli et al. 2023b). To begin exploring intraspecific genetic diversity, we sequenced two Italian *Phy. ampelicida* isolates using short-read technology and analyzed them alongside the available genome of the Spanish PA1 strain (Eichmeier et al. 2022). Despite higher fragmentation, the *de novo* Illumina assemblies of TN2 and LB22.1 showed high completeness (∼99.3% BUSCO), comparable to the PacBio-assembled genome of GW18.1 (**Table 1**). Sequence variants smaller than 50 bp were identified by mapping Illumina reads onto the assembly of GW18.1, while larger structural variants were detected through alignments of whole genome assemblies.

TN2 and LB22.1 contained 89,389 and 89,670 variants relative to GW18.1, respectively (**Supplementary Table 5a**). The majority (90.7 ± 0.0006%) were 1 bp variants, of which 95.5% were SNPs and the remaining 4.5% were 1 bp indels. Variants between 2–10 bp and 11–50 bp accounted for 6.3 ± 0.002% and 2.5 ± 0.01%, respectively. On average, 375 ± 5 variants ranged from 51–250 bp, 89 ± 1 from 251–1000 bp, and 68.5 ± 1.5 from 1001–25,000 bp (**Figure 2A**, **Supplementary Table 5a**). The cumulative variant length averaged 571.6 ± 4.5 kbp per isolate. Single nucleotide variants (including SNPs and 1 bp indels) contributed 14.2 ± 0.4% of the total. Variants in the 11–50 bp, 51–250 bp, and 251–1000 bp ranges each accounted for over 7% of the total length. The longest variants (1001–25,000 bp) made up 58.9 ± 0.4% of the cumulative variant length (**Figure 2B, Supplementary Table 5b**).

**Figure 2.**
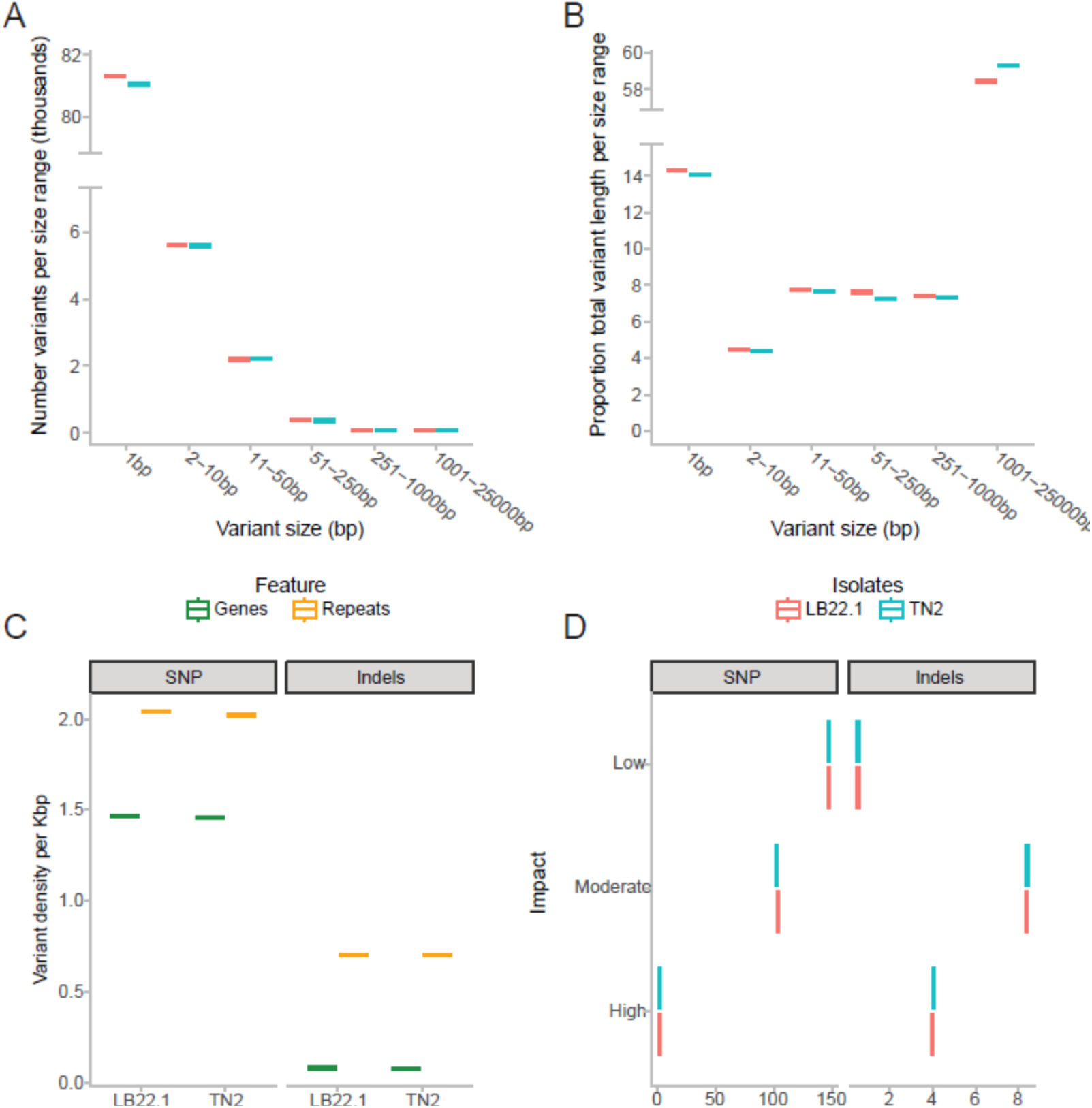
Variant analysis of Phyllosticta ampelicida isolates **A**) Count of variants per size range. **B**) Proportion of total variant length per size range. **C**) Variant density in gene space compared to repeat space. **D**) Number of variants per 100 gene loci and their effect level based on the SNPeff prediction.

The high genetic similarity between the two Italian strains, whose geographical sampling sites are about 150 km apart, likely reflects clonal expansion from a recent common ancestor. Warmer winter temperatures may further suppress sexual reproduction, as observed in *Plasmopara viticola* (Santos et al. 2020), limiting opportunities for genetic recombination. This low diversity could reduce the pathogen’s evolutionary potential, increasing its susceptibility to control measures such as fungicides or host resistance. However, these genomes were assembled using short-read sequencing, which may limit the detection of longer variants.

Variant density in repeat and gene space did not show substantial differences between the Italian strains (**Figure 2C**). As expected, variants were more frequent in repeat regions than in gene space in both isolates (2.7 ± 0.01 *vs.* 1.5 ± 0.003 variants/kbp) (**Supplementary Table 5c**). The average number of variants was significantly lower than expected at chromosome ends (**Supplementary Figure 1**; Chi-squared p-value < 1.03e-3), corresponding to telomeric regions (**Supplementary Table 3**), though other genomic windows emerged as potential evolutionary hot spots (**Supplementary Figure 1**). The lower-than-expected number of variants toward the telomeres could be associated with low mapping quality or reads mapping to multiple regions due to the repetitive nature of telomeric and subtelomeric regions (Treangen et al. 2012). SNPs were more frequent than indels, with marked positional differences. In repeat regions, SNPs were three times more abundant than indels, while in gene space, they were 18 times more frequent (**Figure 2C**). Given that indels are more likely to disrupt gene function, their lower density in gene space than SPNs likely reflects stronger selective constraints on coding regions. This pattern aligns with trends observed in other fungal pathogens, where coding regions are overall conserved, while repeat-rich regions serve as reservoirs for structural variation and adaptive evolution.

Variants were classified based on their predicted impact on coding sequences: low, moderate, or high (e.g., premature stop codons or frameshift mutations). Most affected genes carried low-impact (58.6 ± 0.05%) or moderate-impact (46.8 ± 0.05%) variants, while only 3.4 ± 0.05% were affected by high-impact mutations (**Supplementary Table 5d**). Low-impact and moderate variants were 285 and 12 times more common in SNPs than in indels, respectively. In contrast, high-impact variants were twice as likely to be caused by indels (4.0 ± 0.03 vs. 2.0 ± 0.02 variants per 100 genes) (**Figure 2D, Supplementary Table 5e**).

### Comparative analysis of putative pathogenicity and virulence factor genes

Comparative analysis of family sizes of putative virulence factors was performed between *Phyllosticta* species and plant-pathogenic ascomycetes, focusing on genes potentially associated with pathogenicity and virulence, such as those encoding carbohydrate-active enzymes (CAZymes), cytochrome P450s, biosynthetic gene clusters (BGCs), peroxidases, and cellular transporters. The comparisons included Phyllosticta species with diverse lifestyles: *Phy. citricarpa*, a major citrus pathogen (Timmer et al. 2000); *Phy. capitalensis*, a widespread citrus endophyte and a weak pathogen in other hosts (Silva et al. 2008; Wikee et al. 2013; Cheng et al. 2019); *Phy. citrichinaensis*, a minor citrus pathogen in China which exhibits genomic features of both endophytes and pathogens (Buijs et al. 2022); the closely related *Phy. citribraziliensis*, an endophyte found in asymptomatic citrus leaves in Brazil (Glienke et al. 2011). We also included a non-pathogenic ascomycete (*Sa. cerevisiae*) and two wood-rotting basidiomycetes associated with esca (*F. mediterranea* and *St. hirsutum*), another important grapevine disease.

Overall, the abundance of virulence factor genes confirmed that the Italian strains LB22.1 and TN2 are highly similar to each other and more closely related to the Spanish *Phy. ampelicida* PA1 strain than to the German strain GW18.1. At the interspecific level, *Phy. ampelicida* appears more similar to *Phy. capitalensis* than to *Phy. citrichinaensis*, *Phy. citribraziliensis*, or *Phy. citricarpa* (Sui et al. 2023). Additionally, certain putative virulence factors, such as GH16, AA1, and some secreted CAZymes, were slightly more abundant in *Phy. ampelicida* genomes compared to other Phyllosticta species. However, most other virulence factors analyzed showed similar or lower abundance in *Phy. ampelicida* relative to the rest (**Figure 3**).

**Figure 3.**
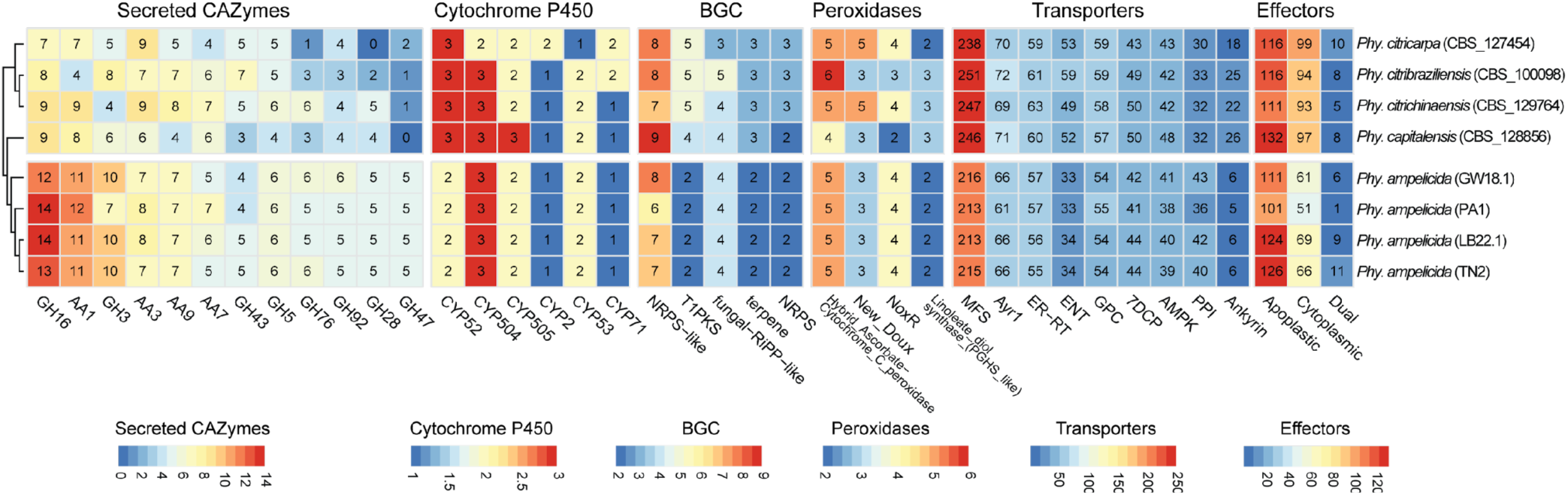
Number of protein-coding genes annotated as secreted CAZymes, P450s, secondary metabolism, peroxidases, transporters, and effectorP prediction. The heatmap includes only the annotations with the highest number of genes across Phyllosticta genomes. Overrepresented (yellow to red) and underrepresented (yellow to blue) domains are depicted as Z-scores for each family.

### Repertoire of carbohydrate-active enzymes

Plant cell wall polysaccharides play a dual role in plant-pathogen interactions, serving as both barriers and nutrient sources (Cantu et al. 2008). Carbohydrate-active enzymes (CAZymes) that target the plant cell walls facilitate these interactions by degrading cellulose, hemicellulose, pectin, and other components, enabling host invasion and nutrient acquisition (Bruno et al. 2006; Ospina-Giraldo et al. 2010; Kubicek et al. 2014; Barrett et al. 2020). A dbCAN3 analysis identified 314 putative CAZyme genes in *Phy. ampelicida*, ∼3% of the total number of genes in the genome (**Table 2**). This is consistent with other *Phyllosticta* species and dothideomycetes, which typically encode 300–400 CAZyme genes (Saier et al. 2006; Ohm et al. 2012; Haridas et al. 2020; Buijs et al. 2021). The largest CAZyme families include glycoside hydrolases (GH, 161 genes), glycosyltransferases (GT, 71 genes), and auxiliary activities (AA, 59 genes). Smaller families include carbohydrate-binding modules (CBM, 16 genes), carbohydrate esterases (CE, 16 genes), and polysaccharide lyases (PL, 7 genes) (**Supplementary Figure 2**).

Among *Phyllosticta* species, *Phy. ampelicida* possesses the smallest set of genes encoding CAZymes, particularly the CBMs, with only 16 genes compared to 34 in *Phy. citribraziliensis* and 45 in *Phy. capitalensis*. CBMs facilitate carbohydrate recognition and enhance enzymatic activity (Gilbert et al. 2013; Várnai et al. 2014). Similarly, *Phy. ampelicida* has fewer GHs (161 vs. ∼177 in other *Phyllosticta* species) and GTs (71 vs. ∼81). Despite these differences, the CAZymes repertoire remains similar across *Phyllosticta* species regardless of their lifestyle (Wang et al. 2020; Buijs et al. 2021). Genome-based prediction of the *Phy. ampelicida* CAZyme secretome, an additional indicator of cell wall-degrading enzyme activity with potential implications during pathogen-plant host interactions, suggests that 50– 85% of CAZyme proteins are secreted, except for glycosyltransferases (GTs) (**Table 3**; **Figure 3, Supplementary Table 8**). The most abundant secreted families target hemicellulose (GH16), cellulose (GH3), and lignin (AA1 laccases). Other major families include AA3 (supporting lignocellulose degradation) and copper-dependent AA9 proteins (Van Den Brink and De Vries 2011; Moses et al. 2016; Sützl et al. 2018; Drula et al. 2022).

**Table 3.** Putative Secreted CAZymes in *Phy. ampelicida* classified by family.

| CAZymes | Total | Secreted | % |
| --- | --- | --- | --- |
| AA | 59 | 39 | 66.10 |
| CBM | 16 | 8 | 50.00 |
| CE | 16 | 11 | 68.75 |
| GH | 161 | 106 | 65.84 |
| GT | 71 | 7 | 9.86 |
| PL | 7 | 6 | 85.71 |

Interestingly, *Phy. ampelicida* has the highest number of predicted secreted CAZymes (∼180) among *Phyllosticta* species, surpassing *Phy. citricarpa* (109) and *Phy. citrichinaensis* (158) (**Figure 3**). For instance, it possesses 14 secreted GH16 proteins—twice as many as *Phy. citricarpa*. Despite this, intra-species variation is minimal, with >90% of secreted CAZyme families showing no differences across isolates. However, the Spanish PA1 isolate exhibits notable reductions in GH3 genes (7 vs. 10 in other isolates) and AA11 lytic polysaccharide monooxygenases (2 vs. 4).

Beyond *Phyllosticta*, pathogenic ascomycetes exhibit a larger CAZyme repertoire, with 114 CAZyme families in *Phaeoacremonium minimum* and 89 in *Elsinoe ampelina*, compared to 56–69 in *Phyllosticta*. These species are enriched in AA7, AA9, and AA3 auxiliary enzymes and GH families targeting pectin (GH43, GH28), hemicellulose (GH16), cellulose (GH3), and chitin (GH18) (Chen et al. 2020; **Figure 4**). The largest families include CBM1 (cellulose-binding) and CE5 (cutinases targeting the plant cuticle). In contrast, *Sa. cerevisiae* has few CAZymes, while wood-decaying basidiomycetes (*F. mediterranea*, *St. hirsutum*) have CAZymes numbers comparable to pathogenic ascomycetes.

**Figure 4.**
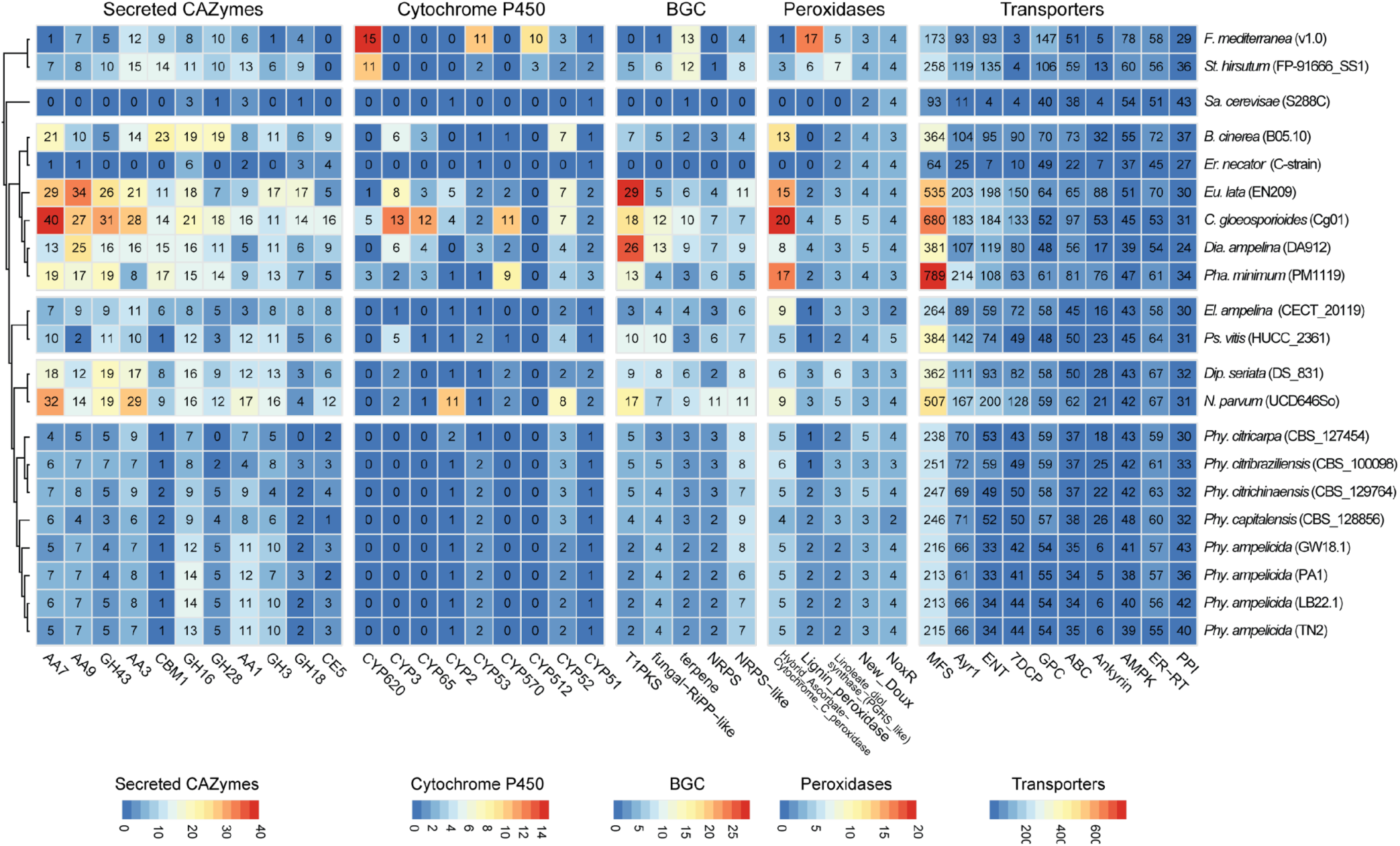
Number of protein-coding genes annotated as secreted CAZymes, P450s, secondary metabolism, peroxidases and transporters. The heatmap includes only the annotations with the highest number of genes across all genomes. Overrepresented (yellow to red) and underrepresented (yellow to blue) domains are depicted as Z-scores for each family.

### Inference of trophic lifestyle based on CAZyme repertoire

CATAstrophy (Hane et al. 2020) categorizes species based on their predicted ecological roles through comparative genomic studies of CAZymes, providing a detailed understanding of their trophic strategies. This classification identifies four ecological groups: monomertrophs, which metabolize simple sugars and include biotrophs and symbionts (both haustorial and non-haustorial species); polymerotrophs, which metabolize complex sugars and encompass necrotrophs, further divided into narrow and broad host-range categories, with the latter requiring a richer CAZyme repertoire for adaptation to multiple hosts; mesotrophs, corresponding to hemibiotrophs, which are further distinguished as intracellular or extracellular based on their adaptation to host invasion, linked to their ability to feed through appressoria-like structures; and vasculartrophs, which are wilt-like pathogens lacking a specific trophic lifestyle but possessing a CAZyme repertoire more similar to broad host-range polymerotrophs.

Species within Phyllostictaceae span multiple trophic classifications, with *Phy. ampelicida* showing an intermediate profile between monomertrophy (1.0), extracellular-mesotrophy (0.89), and saprotrophy (0.81), suggesting a mixed trophic strategy (**Supplementary Table 9**). The number of predicted secreted effectors is consistent with an extracellular-mesotrophic lifestyle, as also supported by its CAZyme profile. However, the boundary between mesotrophs (typically hemibiotrophs) and monomertrophs (biotrophs) remains unclear (De Silva et al. 2016, Précigout et al. 2020). *Phyllosticta ampelicida* has been traditionally classified as a hemibiotroph, exhibiting biotrophic-like features such as a prolonged latent phase exceeding 21 days, followed by a necrotrophic phase characterized by tissue damage and symptom development (Bettinelli et al. 2023a).

### Cytochrome P450 monooxygenases

Cytochrome P450 monooxygenases (CYPs) are a diverse superfamily of heme-containing enzymes involved in the metabolism of endogenous and xenobiotic compounds. In fungi, they play a crucial role in adaptation to ecological niches by contributing to secondary metabolite biosynthesis, nutrient utilization, pathogenesis, resistance to drug and oxidative stress and plant immune responses inhibition (Črešnar and Petrič 2011; Chen et al. 2014a; Durairaj et al. 2016; Fu et al. 2025). In *Phy. ampelicida*, the CYP repertoire is conserved across isolates, with all sharing an identical repertoire of 17 genes across 12 families. Similarly, *Phyllosticta* species exhibit minimal variation, with CYP gene counts ranging from 17 in *Phy. ampelicida* to 19 in *Phy. capitalensis*, *Phy. citribraziliensis*, and *Phy. citrichinaensis*. All species share the same 12 families, except *Phy. citrichinaensis*, which has an additional CYP83 family (**Supplementary Table 8**).

The largest CYP families in *Phyllosticta* include CYP504, an enzyme involved in phenylacetate catabolism (Mingot et al. 1999); CYP52, which aids in alkane and fatty acid assimilation (Ortiz-Álvarez et al. 2020); CYP505, responsible for fatty acid oxidation (Nakayama et al. 1996); and CYP53, which plays a role in detoxification (Moktali et al. 2012; Akapo et al. 2019). Each family contains one to three genes, with little variation among species. However, *Phy. citricarpa*, the most phylogenetically distant species, exhibits distinct differences, possessing an additional CYP2 gene while lacking one gene each in CYP53 and CYP504. Additionally, *Phy. citricarpa* and *Phy. citribraziliensis* share an extra CYP71 gene.

*Phyllosticta* species, along with *El. ampelina*, have the smallest number of CYPs and the fewest identified families, in contrast, Colletotrichum *gloeosporioides* has the largest number of CYPs, with 121 genes spanning 35 families (**Figure 4**). Other pathogenic species exhibit an intermediate number of CYPs, ranging from 31 genes in *Diplodia seriata* and Pseudocercospora *vitis* to 58 genes in *Eutypa lata*. Many CYP families in *Phyllosticta*, such as CYP51 and CYP61, are essential for ergosterol biosynthesis and cell wall integrity (Črešnar and Petrič 2011; Chen et al. 2014b).

### Peroxidases

Peroxidases, a group of oxidoreductases including NAD(P)H oxidase, catalase, and lignin peroxidase, play key roles in lignin degradation, detoxification, and reactive oxygen species (ROS) regulation. These enzymes contribute to carbon recycling and fungal pathogenicity (Choi et al. 2014, Mir et al. 2015, Jia et al. 2023). *Phyllosticta ampelicida* possesses 35 peroxidase genes (34 in strain PA1), distributed across 16 families, each containing one to five genes. Intraspecies variation is minimal, occurring only in the Spanish PA1 strain, which differs from other *Phy. ampelicida* strains and *Phyllosticta* species by a reduced set of atypical 2-cysteine peroxiredoxins. PA1 has only one type II and type V gene instead of two, and it lacks the single type Q and BCP genes typically present in the genus. However, it uniquely possesses the respiratory burst oxidase homolog-type NADPH oxidase gene, which is found in all *Phyllosticta* species but absent in other *Phy. ampelicida* strains.

Among *Phyllosticta* species, the number of peroxidase genes ranges from 33 in *Phy. capitalensis* to 38 in *Phy. citrichinaensis*, with 16 families identified in each species. The largest is the hybrid ascorbate-cytochrome C peroxidase family, averaging 5±0.3 genes per species. The most notable differences are found in the NADPH oxidase (Nox) enzyme family, which synthesizes ROS for cell defense and signaling (O’Brien et al. 2012). While NoxA and NoxB are conserved across the genus, *Phy. ampelicida* is the only species to possess a NoxC gene, a rare subfamily with an unclear biological function (Takemoto and Scott 2023). Additionally, the two endophytic species, *Phy. citribraziliensis* and *Phy. capitalensis*, have a reduced set of NoxR regulatory genes.

Compared to other grapevine pathogens, *Phyllosticta* species have the fewest peroxidase genes, except for *Erysiphe necator* (22 genes). Other pathogens possess larger repertoires, ranging from 47 in *El. ampelina* to 58 in *N. parvum*, with *C. gloeosporioides* having the most (71 genes; **Figure 4**). Across these species, heme peroxidases are the predominant type, with the hybrid ascorbate-cytochrome C peroxidase family being particularly abundant. These enzyme families are quite large in pathogens such as *C. gloeosporioides*, *Pha. minimum*, *Eu. lata*, and *Botrytis cinerea*, where their numbers range from 13 to 20 genes.

### Biosynthetic gene clusters and secondary metabolism potential

Fungal secondary metabolites are essential for growth, development, pathogenesis, nutrient acquisition, and ecological interactions (Keller 2019; Yang et al. 2024). In fungal genomes, genes encoding enzymes involved in biosynthesis, modification, and transport of secondary metabolites are organized into biosynthetic gene clusters (BGCs), which typically include core biosynthetic genes, tailoring enzymes, regulatory transcription factors, and transporters (Brakhage 2013; Keller 2019). *Phyllosticta ampelicida* strains share a small core set of BGCs. Italian strains TN2 and LB22.1 have 20 BGCs each, while the Spanish strain PA1, which lacks the isocyanide cluster and has only six NRPS-like clusters, has the fewest (18). The GW18.1 strain, with eight NRPS-like clusters, has the highest number (21) (**Supplementary Table 8**). Within *Phyllosticta*, *Phy. ampelicida* has the fewest BGCs, whereas *Phy. citrichinaensis* and *Phy. citribraziliensis* contain 25 and 28 clusters, respectively. *Phyllosticta ampelicida* lacks the NRP-metallophore BGC and has the smallest type I polyketide synthase (T1PKS) cluster, with an overall moderate reduction in BGCs. Despite this, a minimal core set of BGCs is shared within the genus.

Secondary metabolite diversity varies widely among grapevine pathogens (**Supplementary Table 8**). *Diaporthe ampelina* has the highest number of BGCs (81), followed by *Eu. lata* (69) and *N. parvum* (67). In contrast, *Phyllosticta*, *El. ampelina* (23), and *B. cinerea* (25) have fewer. Half of the BGCs are species-specific, while others, such as fungal-RiPP clusters, are found in only two species. Common grapevine pathogen BGCs (T3PKS, indole, NRPS|T1PKS-like) are absent in *Phyllosticta*, while beta-lactone, isocyanide, and NRP-metallophore|NRPS clusters are common in *Phyllosticta* but rare in other pathogens.

### Transporters

Membrane transporters contribute to virulence by secreting secondary metabolites and toxins while excluding host-derived antimicrobial compounds (Denny and VanEtten 1983; Denny et al. 1987). Among *Phy. ampelicida* isolates, transporter variation is minimal (**Supplementary Table 8**), with the major facilitator superfamily (MFS) being the largest. At the genus level, *Phy. ampelicida* has the fewest transporters, with the numbers of members being reduced across all major families. Compared to other grapevine pathogens, *Phyllosticta* species exhibit fewer transporters, particularly within the MFS family. Overall, the clade including *Eu. lata, C. gloeosporioides, Dia. ampelina,* and *Pha. minimum* has the highest number of identified virulence factors, while *Phyllosticta* possesses relatively few.

### Putative fungicide targets

*Phyllosticta ampelicida* exhibits insensitivity to Cidely, a fungicide targeting grapevine powdery mildew. Cidely combines Cyflufenamid (FRAC Code U6, unknown mode of action) and Difenoconazole (FRAC Group 3, demethylation inhibitor or sterol biosynthesis inhibitor), which disrupts ergosterol biosynthesis, essential for fungal cell membrane integrity. DMI resistance in fungi often stems from mutations in *CYP51*, encoding cytochrome P450 lanosterol C-14α demethylase. However, the gene Gb01.g2769 in the sequenced *Phy. ampelicida* GW18.1 lacks known fungicide resistance mutations, such as Y137F (Behr et al. 2012), F489L (Mair et al. 2020), M231T (Zhang et al. 2020), S509T (Arnold et al. 2024), Y144F/Y144H (Pereira et al. 2012), I387M (Muellender et al. 2021), and G207A (Wang et al. 2021). This suggests its insensitivity to Cidely may stem from alternative mechanisms, such as enhanced efflux pump activity or increased *CYP51* accumulation (Jones et al., 2014).

### Gene family expansion and contraction

To assess whether differences in putative virulence factor counts among species groups result from accelerated gene family evolution, we conducted a CAFE analysis (Mendes et al. 2021). This approach calculates gene birth and death rates within families to identify lineages with significant changes in gene family size. First, single-copy orthologs were identified in predicted proteins across all genomes and used to build a clock-calibrated tree (**Figure 5**), calibrated with ascomycetes crown age (588 Mya) and dothideomycetes divergence (350 Mya). Second, all predicted proteins were clustered into families, and gene family sizes were calculated. Both datasets were used to run the CAFE pipeline. *Phyllosticta* species exhibit a low expansion/contraction ratio, with only three significantly expanded and 165 significantly contracted gene families (**Figure 5**). Specifically, the *Phy. ampelicida* lineage shows an overall contraction, with 10 expanded and 65 contracted gene families. In contrast, highly pathogenic species like *Eu lata, Pha. minimum,* and *N. parvum* show at least 50% more expanding than contracting gene families. This difference may relate to *Phy. ampelicida* extended latent period, resembling biotrophic organisms (Bettinelli et al, 2023a), while the necrotrophic *Eu. lata, Pha. minimum,* and *N. parvum* exhibits high virulence. Another factor could be *Phy. ampelicida* narrow host range, restricted to Vitaceae (Szabó et al. 2023), compared to the broad host range of *Eu. lata, Pha. minimum,* and *N. parvum* (Garcia et al. 2021).

**Figure 5.**
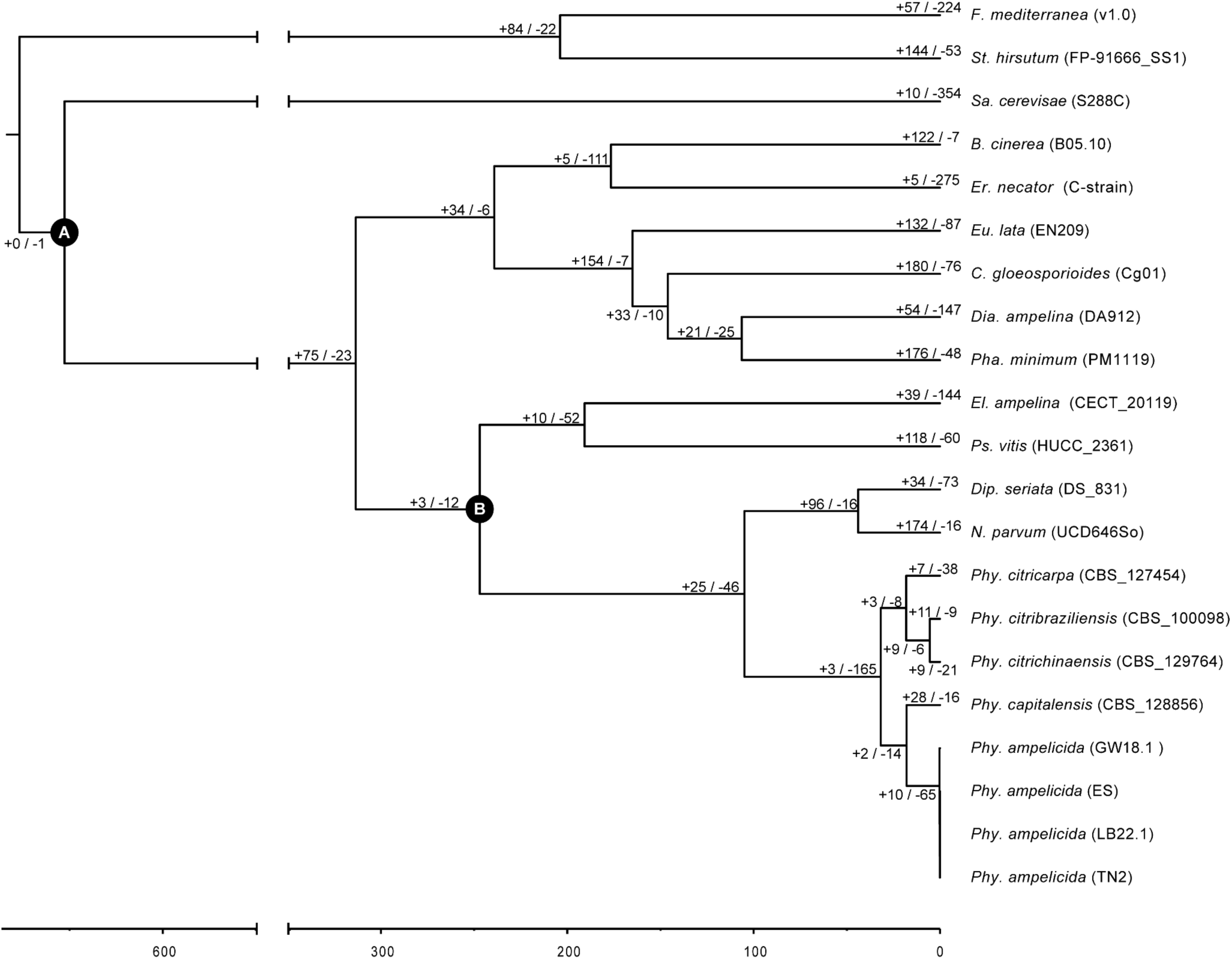
Clock-calibrated phylogenetic tree with estimated times of divergence in million years ago and numbers of expanded and contracted gene families. Clock-calibrated phylogenetic tree with analysis of gene family expansion and contraction. The length of the branches represents divergence times in million years. Calibration point (**A**) at ascomycete crown. Calibration point (**B**) at the dothideomycetes divergence. Positive and negative numbers represent families under expansion and contraction, respectively, as determined by gene family evolution analysis CAFE.

Genes from significantly expanded or contracted families were analyzed for functional enrichment (**Figure 6**). In *Phy. citricarpa* and *Phy. citribraziliensis*, expansion is enriched for APC (amino acid-polyamine-organocation) superfamily transporters (2.A.39, 2.A.21, 2.A.3), which facilitate ion co- and counter-transport with amino acids, likely contributing to nutrient acquisition and other cellular processes (Jack et al. 2000). The 9.B.12 transporter family, linked to salt tolerance, is enriched in *Phy. citrichinaensis* expanding families. In *Sa. cerevisiae*, deletion of genes in this family increases NaCl sensitivity (Navarre 2000). In *Phy. capitalensis*, α-Type channels (1.A) are enriched in contracted families, while in *Phy. ampelicida*, they are enriched in expanding families. These ubiquitous transporters facilitate passive solute movement (Saier 2000). This is consistent with the initial and elongated biotrophic phase of *Phy. ampelicida* in the development of black rot in grapevine, where the fungi limits its interaction with few superficial cells (Kuo and Hoch 1996a; Ullrich et al. 2009). Expanding this kind of transporters possibly helps the fungi with a slow but steady development without spending much resources in the transport of nutrients. Additionally, the contracted families of *Phy. ampelicida* show an enrichment for CAZymes, including glycoside hydrolases (GH78, GH2), carbohydrate-binding modules (CBM63), and auxiliary activity enzymes (AA3, AA9). These genes function in rhamnosidase and galactosidase activity, lignocellulose binding and degradation, and lytic polysaccharide monooxygenase activity, all associated with fungal degradation of plant cell walls. The contraction of these functions in *Phy. ampelicida* suggests a reduced ability to degrade plant material, particularly mature tissues with thickened cuticles and cell walls (Kuo & Hoch, 1996), especially when compared with highly pathogenic species like *Eu. lata,* and *N. parvum* where this groups of functions seem are enriched in the families under expansion (Morales-Cruz et al. 2015; Garcia et al. 2021).

**Figure 6.**
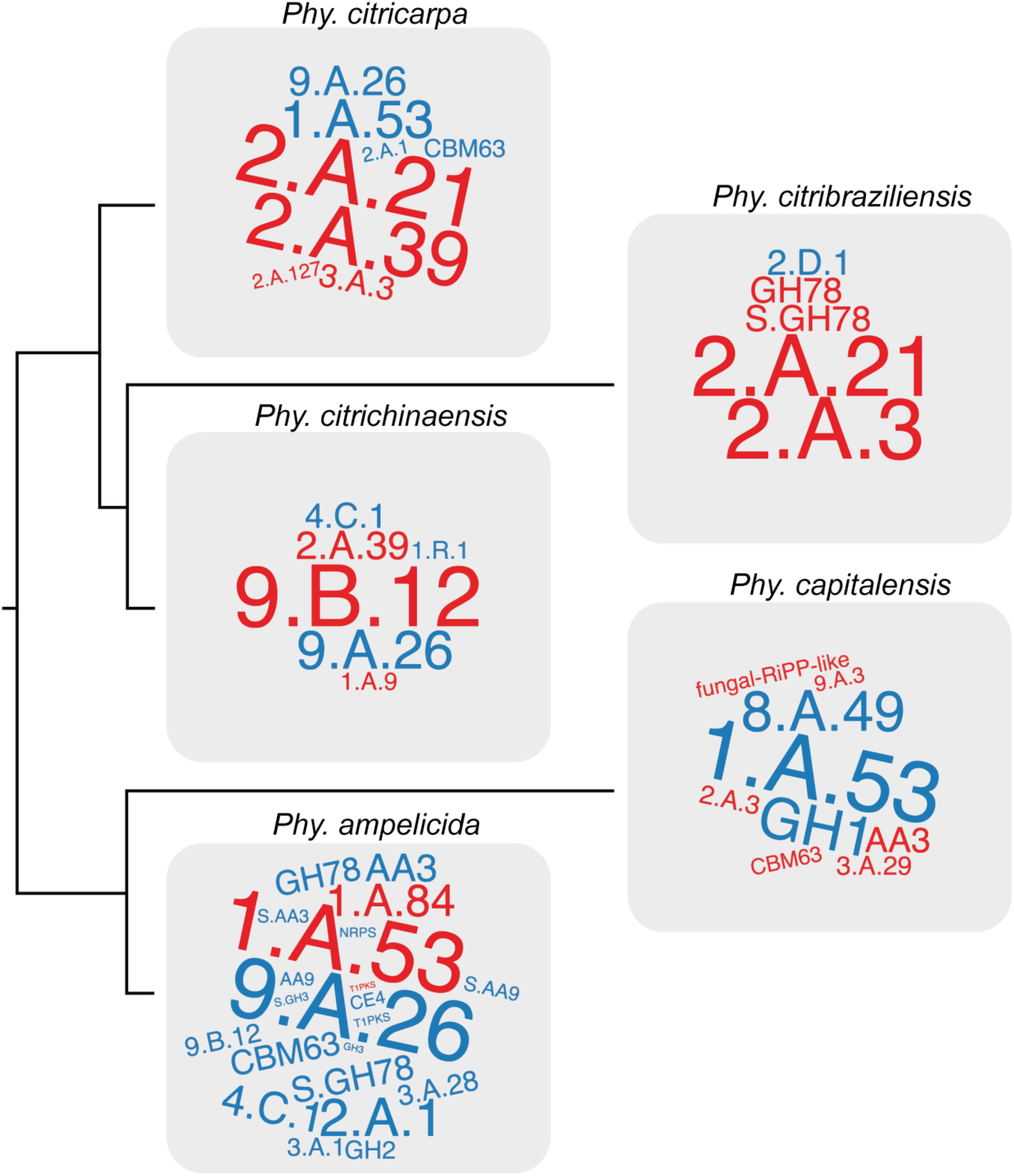
Word clouds representing the genes enriched in the rapidly evolving families of the Phyllosticta species presented in this work. The word size represents the enrichment’s strength based on the P value. The red color represents the enriched genes within expanding families, and the blue represents enriched genes within contracting families.

## Supplementary Materials

Supplemental material available at:

## Author Contributions

Conceptualization: MC, PB, JG, SV, SM, DC

Formal analysis: MC, PB, JG

Investigation: MC, PB, JG, GM

Resources: MC, PB, JG, GM, ST, LH, SV, SM, DC

Data curation: JG

Writing—original draft preparation: MC, PB; DC

Writing—review and editing: MC, PB, JG, ST, LH, SV, SM, DC

Visualization: MC, PB, JG

Supervision: MC, SV, SM, DC

Project administration: MC, SV, SM, DC

Funding acquisition: SM, DC

All authors have read and agreed to the published version of the manuscript.

## Funding

CM, SLT, GM, SM were supported by Regione Lombardia (project 33 New Defense Strategies Against Black Rot in Grapevines: A Threat to Lombard Viticulture CUPG44I20000890003). DC was partially supported by the Ray Rossi Endowment in Viticulture and Enology and by the California Department of Food and Agriculture, California Fruit Tree, Nut Tree, and Grapevine Improvement Advisory Board (Grant# 20-1062-000-SA; 21-0427-000-SA; 22-1588-000-SA; 23-0706-000-SA). PB was supported by the Agritech National Research Center and received funding from the European Union Next-Generation EU (PIANO NAZIONALE DI RIPRESA E RESILIENZA (PNRR)—MISSIONE 4 COMPONENTE 2, INVESTIMENTO 1.4—D.D. 1032 17/06/2022, CN00000022).

## Data Availability

The sequencing data and genome assemblies for this project are available at NCBI (https://www.ncbi.nlm.nih.gov/bioproject/PRJNA1231451). The genome assemblies and gene models produced in this study are publicly available at Zenodo (https://zenodo.org/records/15264932). Dedicated genome browsers and BLAST tools are available at https://www.grapegenomics.com.

## Acknowledgments

We thank Drs. Rosa Figueroa-Balderas, Andrea Minio, and Noé Cochetel for their technical support and the UC Davis DNA Technologies Core for sequencing assistance.

## Conflicts of Interest

The authors declare no conflict of interest.

